# Spatial relationships of intra-lesion heterogeneity in *Mycobacterium tuberculosis* microenvironment, replication status, and drug efficacy

**DOI:** 10.1101/2021.11.08.467819

**Authors:** Richard C. Lavin, Shumin Tan

## Abstract

A hallmark of *Mycobacterium tuberculosis* (Mtb) infection with critical impact on disease development and outcome is the marked heterogeneity that exists, spanning differences in lesion types to changes in microenvironment as the infection progresses^1–7^. A mechanistic understanding of how this heterogeneity affects Mtb growth and treatment efficacy necessitates single bacterium-level studies in the context of intact host tissue architecture; however, such an evaluation has been technically challenging. Here, we exploit fluorescent reporter Mtb strains and the C3HeB/FeJ murine model in an integrated imaging approach to study microenvironment heterogeneity within a single lesion *in situ*, and analyze how these differences relate to non-uniformity in Mtb replication state, activity, and drug efficacy. We show that the pH and chloride environments differ spatially in a caseous necrotic lesion, with increased acidity and chloride levels in the lesion cuff versus the necrotic core. Conversely, a higher percentage of Mtb in the necrotic core versus the lesion cuff were in an actively replicating state, and correspondingly active in transcription and translation. Finally, examination of three first-line anti-tubercular drugs showed that efficacy of isoniazid was strikingly poor against bacteria in the lesion cuff. Our study reveals spatial relationships of intra-lesion heterogeneity, sheds light on important considerations in the development of anti-tubercular treatment strategies, and establishes a foundational framework for Mtb infection heterogeneity analysis at the single cell level *in situ*.

## MAIN TEXT

The ability to effectively treat *Mycobacterium tuberculosis* (Mtb) is significantly impeded by the marked heterogeneity of the infection across multiple levels, including non-uniformity in local microenvironments^1–6^. This heterogeneity extends not just between lesions but within a single lesion; for example, matrix-assisted laser desorption/ionization mass spectrometry studies have demonstrated variation in drug penetration into caseous necrotic lesions^8^. Further, pH measurements of liquefying caseum obtained from caseous necrotic lesions in both C3HeB/FeJ mice and guinea pigs have indicated the neutral pH of this material^8,9^, which has been contrasted with the slightly acidic pH of the macrophage intraphagosomal environment that is a major niche of the bacterium^10–12^. Of note, the first-line anti-tubercular drug pyrazinamide (PZA) shows increased efficacy in acidic conditions^13–15^. The neutral pH of the caseous material has thus been implicated as a contributing reason for the lack of PZA efficacy sometimes observed in C3HeB/FeJ mice where caseous necrotic lesion types are formed, versus the uniform efficacy observed in BALB/c mice, which do not form caseous necrotic lesions^8,9^.

The critical impact of within-host heterogeneity on infection and treatment outcome dictates the need to functionally characterize Mtb infection *in vivo* at the single bacterium level, within spatial tissue context. However, the technical hurdles associated with accomplishing such studies has meant a continued dearth in our knowledge of what Mtb actually “sees” during infection at the single bacterium level, and how this may be non-uniform within a host or lesion. In addition, we lack vital information regarding how bacterial growth status may vary in different sublocations within a host. To overcome these challenges and address these key questions, we sought to develop an integrated imaging approach that enables analysis of single bacteria in individual lesions within an infected lung. To establish this approach, we infected C3HeB/FeJ mice with Mtb constitutively expressing mCherry and harvested the lungs 6 weeks post-infection. The utility of C3HeB/FeJ mice as a Mtb infection model has been increasingly appreciated due to its formation of a range of lesion types including caseous necrotic lesions, and it is now frequently used in anti-tubercular drug studies^8,9,16–21^. By employing a broad xy-plane tiled imaging approach coupled with antibody staining against host markers, we were able to distinguish the three lesion types previously described via histological studies in the C3HeB/FeJ murine Mtb infection model (Supplementary Fig. 1)^4,7^. Of particular interest here, highly structured type I caseous necrotic lesions were distinguished by the fibrous collagen I-rich cuff containing CD68-positive macrophages that rings the caseous necrotic core (Supplementary Fig. 1a). Very rare neutrophil-dominant type II lesions were discriminated by Ly6G staining for neutrophils (Supplementary Fig. 1b), while macrophage-rich type III lesions were differentiated with CD68 staining of macrophages (Supplementary Fig. 1c). Broad xy-plane tiled imaging provides the breadth required to capture large type I lesions in their entirety (Supplementary Fig. 1a), with subsequent targeted 3-dimensional imaging and reconstruction enabling single cell-resolution visualization of the fluorescent bacteria (Supplementary Fig. 1, insets), setting the stage for analysis of intra-lesion sublocation environment and Mtb replication status. We focus here on type I caseous necrotic lesions, due to its association with heterogeneity in drug penetration and Mtb drug response^8,9,16–18,22^, and its highly structured nature.

We first sought to directly visualize differences in the pH and chloride (Cl^−^) microenvironment within type I caseous necrotic lesion sublocations, given the reported neutral pH of liquifying caseum^8,9^, and our previous work demonstrating the synergistic response of Mtb to pH and Cl^−^, which are linked cues during macrophage phagosomal maturation^12^. To do so, we infected C3HeB/FeJ mice with our pH/Cl^−^-responsive fluorescent reporter Mtb strain (Erdman *rv2390c’*::GFP, *smyc’*::mCherry) that fluoresces green upon bacterial exposure to acidic pH and/or high Cl^−^levels in the local environment, and expresses mCherry constitutively for visualization of all bacteria irrespective of local environment^12,23^. Comparison of Mtb present in the lesion cuff (predominantly present within macrophages) versus those present in the necrotic lesion core 6-8 weeks post-infection showed that the *rv2390c’*::GFP reporter signal was significantly higher in the bacteria present in the lesion cuff (Figs. 1a-1c), even as non-uniformity in reporter signal within each sublocation remained notable (Fig. 1c). Still more strikingly, binning the data for each lesion into different *rv2390c’*::GFP reporter signal ranges demonstrated the opposite distributions in physiological cues experienced by the two bacterial populations (Fig. 1d). Specifically, most Mtb present in the lesion cuff expressed high levels of *rv2390c’*::GFP reporter fluorescence (Figs. 1b-1d), indicative of an environment with more acidic pH and/or higher [Cl^−^], and in accord with the environment that would be expected in macrophage phagosomes^12^. Conversely, a majority of bacteria present in the necrotic core (extracellular Mtb) expressed lower levels of *rv2390c’*::GFP reporter signal (Figs. 1b-1d), indicative of an environment that is at a more neutral pH/has a lower [Cl^−^]. Our findings demonstrate at the single bacterium level within intact tissue that (i) the pH and chloride environment of the caseous necrotic core significantly differs from that experienced by Mtb present within the lesion cuff, and (ii) there remains non-uniformity in the local environment within each sublocation, even in the necrotic core.

**Fig. 1.**
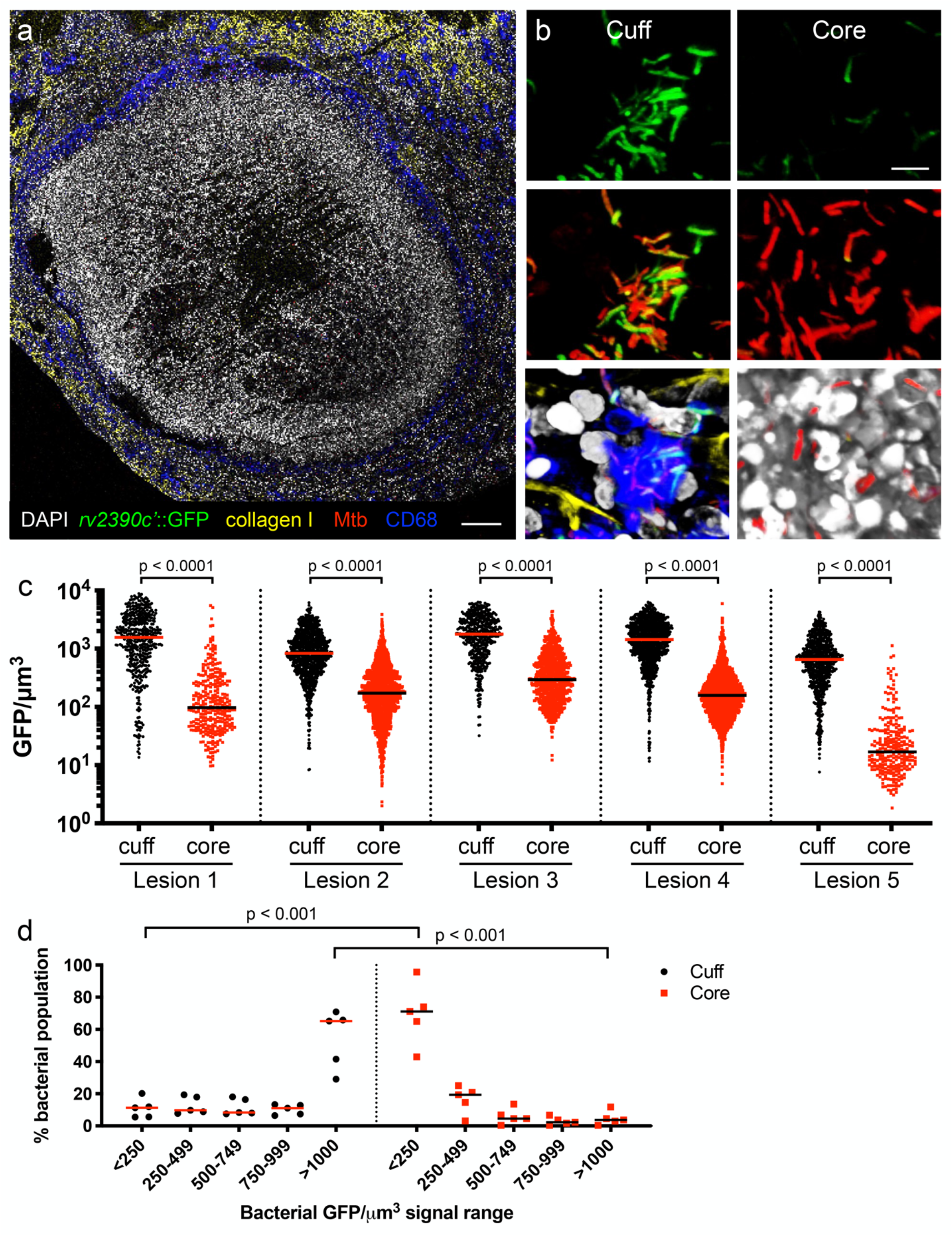
*rv2390c’*::GFP reporter reveals heterogeneity in pH/Cl^−^levels within different sublocations of caseous necrotic lesions. (a and b) Confocal microscopy images of type I caseous necrotic lesions from 6-8 week infection of C3HeB/FeJ mice with Erdman (*rv2390c’*::GFP, *smyc’*::mCherry). Overview image of a lesion (~10 × 11 tiled image) is shown in (a), and representative 3D confocal images from the lesion cuff and core shown in (b). All bacteria are marked in red (*smyc’*::mCherry), reporter signal is shown in green (*rv2390c’*::GFP), nuclei are shown in grayscale (DAPI), collagen I is shown in yellow, and macrophages are shown in blue (CD68). Scale bar 200 μm in (a) and 5 μm in (b). (c) shows GFP/μm^3^ signal for individual bacteria or a group of tightly clustered bacteria, quantified from multiple 3D confocal images at each lesion sublocation (5 different lesions from 5 mice; number of bacteria quantified was respectively 478, 307, 937, 1715, 516, 885, 1078, 1755, 749, and 257 for each lesion sublocation as shown from left to right on the graph). Horizontal lines mark the median value for each sample. p-values were obtained with a Mann-Whitney statistical test. (d) shows data from (c) binned into 5 sub-ranges of GFP/μm^3^ signal. Each point on the graph represents one lesion. Horizontal lines mark the median value for each group. p-values were obtained with a multiple t-test with a Holm-Sidak correction.

To understand how this intra-lesion heterogeneity in microenvironment affects Mtb infection and treatment outcome, we next utilized our previously described single-strand DNA-binding protein (SSB)-GFP replication reporter to determine the replication status of Mtb in sublocations within a lesion. In this reporter strain, the Mtb SSB protein is translationally fused to GFP, driven by the native *ssb* promoter, and Mtb undergoing active DNA replication exhibit green foci, providing a proxy for revealing the replication status of a given bacterium^23,24^. Strikingly, analysis of lung tissue from a 6 week C3HeB/FeJ mice infection with the Erdman (SSB-GFP, *smyc’*::mCherry) reporter Mtb strain showed that a significantly greater percentage of Mtb present in the lesion core possessed SSB-GFP foci, indicative of actively replicating bacteria, versus Mtb present in the lesion cuff (Fig. 2). This difference in overall replication status between Mtb present in the two lesion sublocations fits with the observed differences in local pH/[Cl^−^] (Fig. 1), with a higher percentage of replicating Mtb in the less harsh environment of the necrotic core. The replication state of Mtb can differentially impact drug efficacy and is implicated as a major contributor to the difficulties of successfully treating Mtb infection, as well as to the necessity for a prolonged treatment time course^25–29^. Yet actual demonstration of how Mtb replication status may differ within a single host during infection has been difficult to establish. Our data presented here provides the first direct evidence, to our knowledge, of how Mtb replication status differs not just within a single host but within a single lesion, and reveals how the non-uniformity in intra-lesion Mtb growth status is spatially related to lesion architecture.

**Fig. 2.**
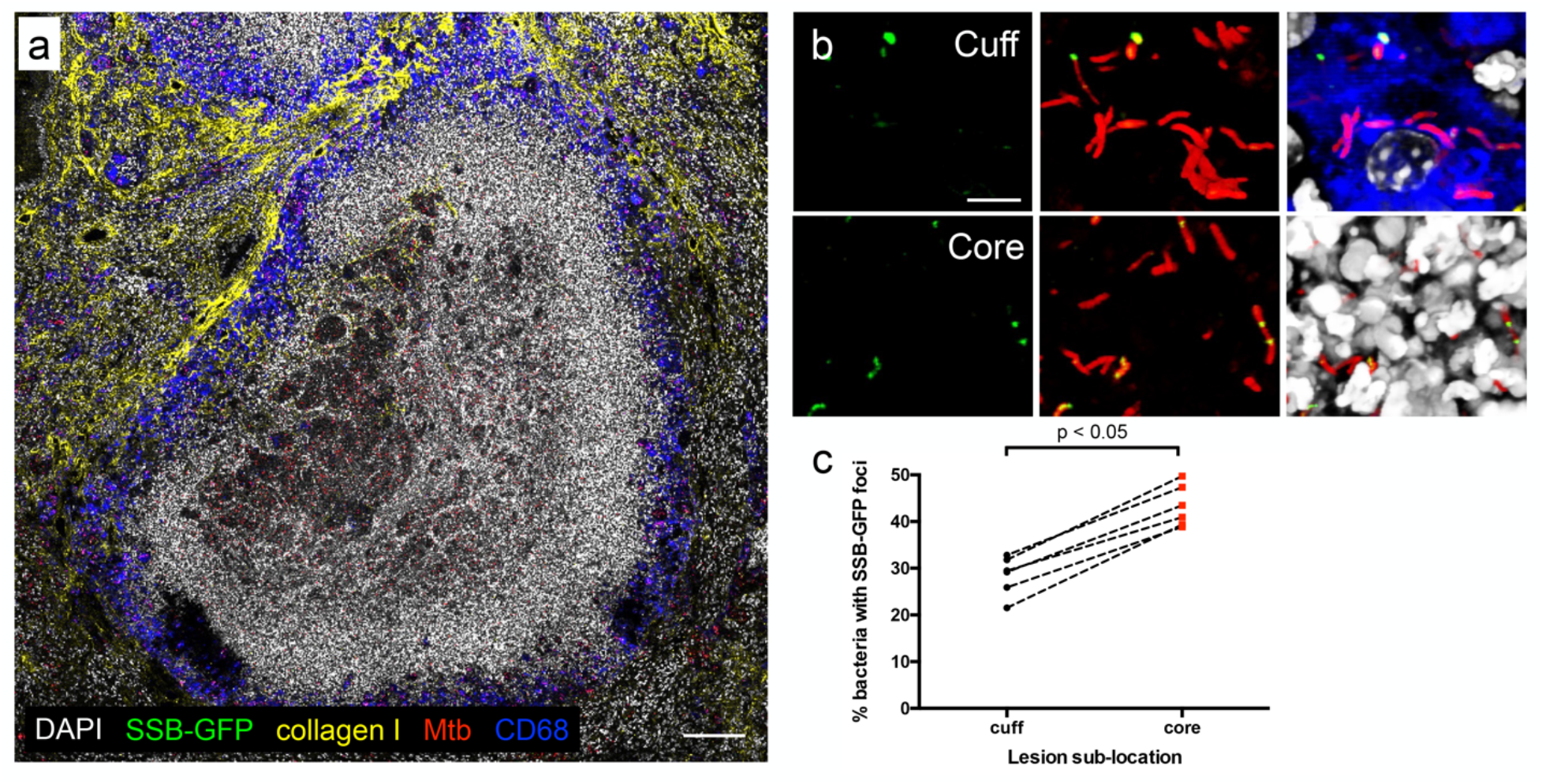
SSB-GFP reporter reveals heterogeneity in bacterial replication within different sublocations of caseous necrotic lesions. (a and b) Confocal microscopy images of type I caseous necrotic lesions from 6 week infection of C3HeB/FeJ mice with Erdman (SSB-GFP, *smyc’*::mCherry). Overview image of a lesion (~5 × 5 tiled image) is shown in (a), and representative 3D confocal images from the lesion cuff and core are shown in (b). All bacteria are marked in red (*smyc’*::mCherry), reporter signal is shown in green (SSB-GFP), nuclei are shown in grayscale (DAPI), collagen I is shown in yellow, and macrophages are shown in blue (CD68). For clarity of foci visualization, SSB-GFP signal is shown in extended focus, overlaid on the 3D images in (b). Scale bar 100 μm in (a) and 5 μm in (b). (c) shows the percentage of Mtb displaying SSB-GFP foci in each lesion sublocation for each quantified lesion measured from multiple 3D confocal images (6 different lesions from 5 mice; number of bacteria quantified in each lesion sublocation [cuff, core] was [678, 475], [564, 545], [370, 446], [478, 1248], [537, 867], and [393, 360]). p-value was obtained using a Wilcoxon matched-pairs signed rank test.

The marked difference in bacterial replication state of Mtb residing in the caseous necrotic lesion core versus cuff prompted us to test a second, independent, approach to analyzing Mtb physiological state *in situ*. A dual fluorescent system where one fluorophore is placed under the control of an inducible promoter and a second spectrally distinct fluorophore is expressed constitutively has been successfully utilized to differentiate between transcriptionally active versus non-active Mtb in cultured macrophages^30–32^. We thus applied this strategy here *in vivo*, exploiting a reporter Mtb strain that carries on the chromosome a tetracycline inducible monomeric Kusabira Orange (mKO) construct, along with a constitutively expressed mCherry (P_606_’::mKO-tetON, *smyc’*::mCherry)^30,33^. As an initial test of the system, we infected C3HeB/FeJ mice with the Erdman (P_606_’::mKO-tetONm *smyc’*::mCherry) reporter Mtb strain for one week, before the provision of drinking water containing 5% sucrose ± 1 mg/ml doxycycline (dox) for one additional week. As shown in Figs. 3a-3c, in this short-term infection, dox treatment resulted in the expected induction of mKO fluorescence in Mtb, with Mtb in the mock-treated mice displaying no mKO fluorescence, reinforcing the suitability of dox induction for *in vivo* Mtb studies^34^. To assess spatial intra-lesion differences in Mtb transcriptional/translational activity, we infected C3HeB/FeJ mice with the Erdman (P_606_’::mKO-tetON, *smyc’*::mCherry) reporter Mtb strain, allowing the infection to establish for 6 weeks prior to a one week exposure of the mice to drinking water containing 5% sucrose + 1 mg/ml dox, and harvesting of lung tissue. A first observation revealed by broad xy-plane imaging was that penetration of dox into the very central core of the caseous necrotic lesion appeared impeded, as no mKO signal could be observed in the bacteria present there (Supplementary Fig. 2). Nonetheless, as mKO signal could be observed within the more peripheral regions of the caseous necrotic core (Supplementary Fig. 2), analysis of the differences in mKO induction in Mtb present in the caseous necrotic core versus cuff was still feasible. Conspicuously, induction of mKO signal was observed in a much greater percentage of Mtb present in the peripheral regions of the caseous necrotic core versus those in the lesion cuff (Figs. 3d and 3e). Additionally, the variation in mKO signal induction within both the cuff and core bacterial populations was significantly greater than that observed in the short-term infection (compare Fig. 3b and 3e). These data are consistent with the differences in bacterial replication status between Mtb present in the caseous necrotic core versus cuff, and further supports and highlights the impact of lesion sublocation on Mtb physiological state.

**Fig. 3.**
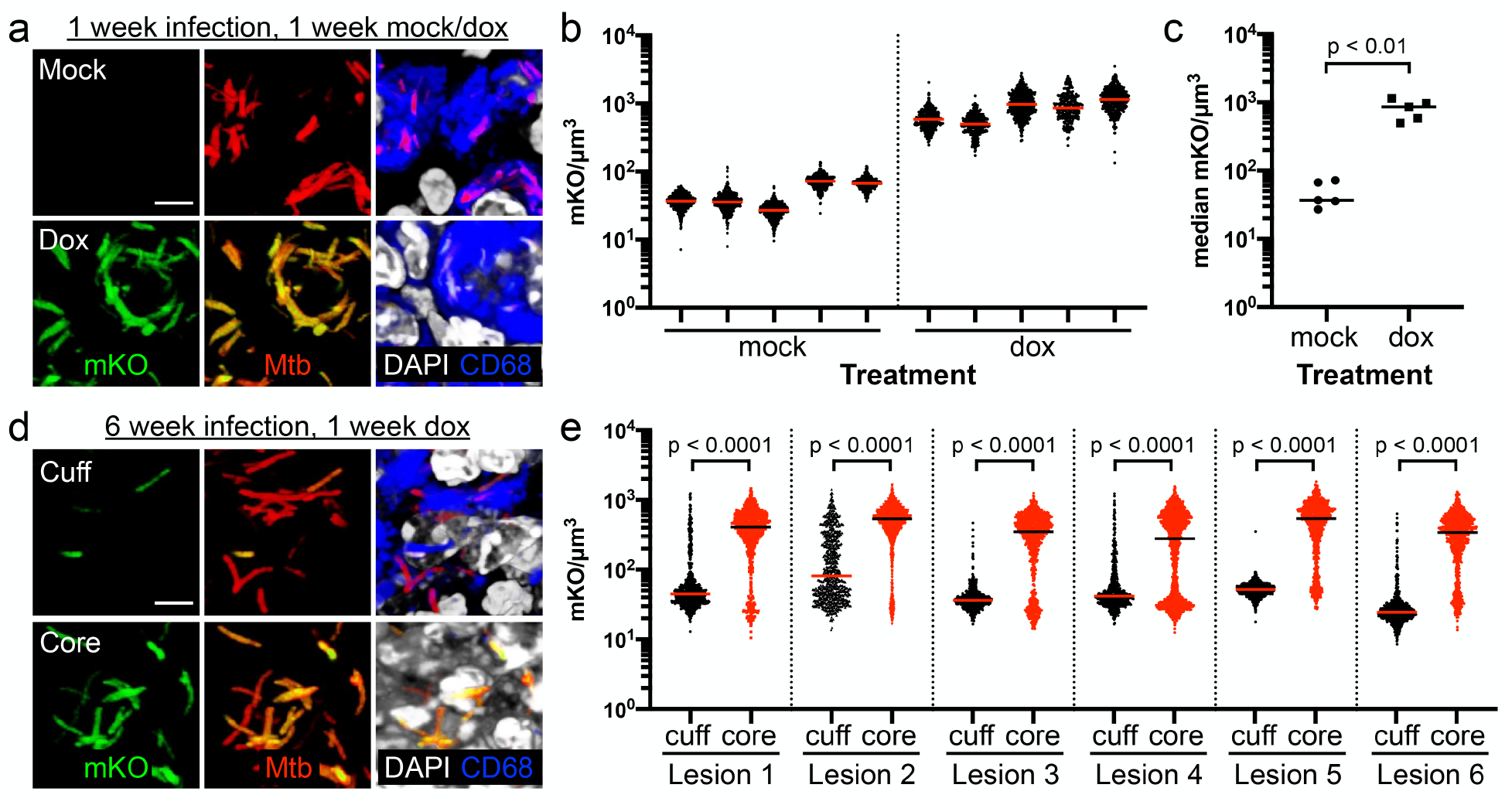
Heterogeneity in bacterial transcriptional/translational activity within different sublocations of caseous necrotic lesions. (a) Representative 3D confocal images from the lesion cuff and core from a 1 week infection of C3HeB/FeJ mice with Erdman (P_606_’::mKO-tetON, *smyc’*::mCherry), followed by 1 week of exposure to drinking water ± 1 mg/ml doxycycline. All bacteria are marked in red (*smyc’*::mCherry), reporter signal is shown in green (P_606_’::mKO-tetON), nuclei are shown in grayscale (DAPI), and macrophages are shown in blue (CD68). Scale bar 5 μm. (b) mKO/μm^3^ signal for individual bacteria or a group of tightly clustered bacteria, quantified from multiple 3D confocal images from infections as performed in (a) (5 different mice/treatment group; number of bacteria quantified from each sample was respectively 440, 424, 604, 379, 436, 355, 330, 545, 257, and 425 as shown from left to right on the graph). Horizontal lines mark the median value for each sample. (c) graphs the medians of each sample shown in (b), with p-value obtained with a Mann-Whitney statistical test. (d) Representative 3D confocal images from the lesion cuff and core from a 6 week infection of C3HeB/FeJ mice with Erdman (P_606_’::mKO-tetON, *smyc’*::mCherry), followed by 1 week of exposure to drinking water + 1 mg/ml doxycycline. All bacteria are marked in red (*smyc’*::mCherry), reporter signal is shown in green (P_606_’::mKO-tetON), nuclei are shown in grayscale (DAPI), and macrophages are shown in blue (CD68). Scale bar 5 μm. (e) mKO/μm^3^ signal for individual bacteria or a group of tightly clustered bacteria, quantified from multiple 3D confocal images at each lesion sublocation (6 different lesions from 5 mice; number of bacteria quantified was respectively 612, 745, 728, 1802, 488, 913, 636, 1615, 1527, 1058, 599, and 921, for each lesion sublocation as shown from left to right on the graph). Horizontal lines mark the median value for each sample. p-values were obtained with a Mann-Whitney statistical test.

The observed differences in pH/Cl^−^environment and in bacterial replication status and transcriptional/translational activity between Mtb residing in the lesion core versus cuff raised the question of whether anti-tubercular drugs may have differential efficacy even within a single caseous necrotic lesion, separate from lesion penetration issues. To address this question, we infected C3HeB/FeJ mice for 6 weeks with the Erdman (SSB-GFP, *smyc’*::mCherry) reporter Mtb strain, before starting treatment via oral gavage with isoniazid (INH), PZA, or rifampicin (RIF) for two weeks. A control set of infected mice were mock-treated with sterile water. Prominently, consistent with the better efficacy of INH against actively replicating bacteria^35–37^, INH treatment was significantly more effective against Mtb present in the caseous necrotic core versus the cuff, with a decrease in the percentage of Mtb with SSB-GFP foci compared to the mock-treated mice in the lesion core (Fig. 4). In contrast, no significant difference in the percentage of Mtb with SSB-GFP foci was observed between the INH and mock-treated groups in the lesion cuff (Fig. 4).

**Fig. 4.**
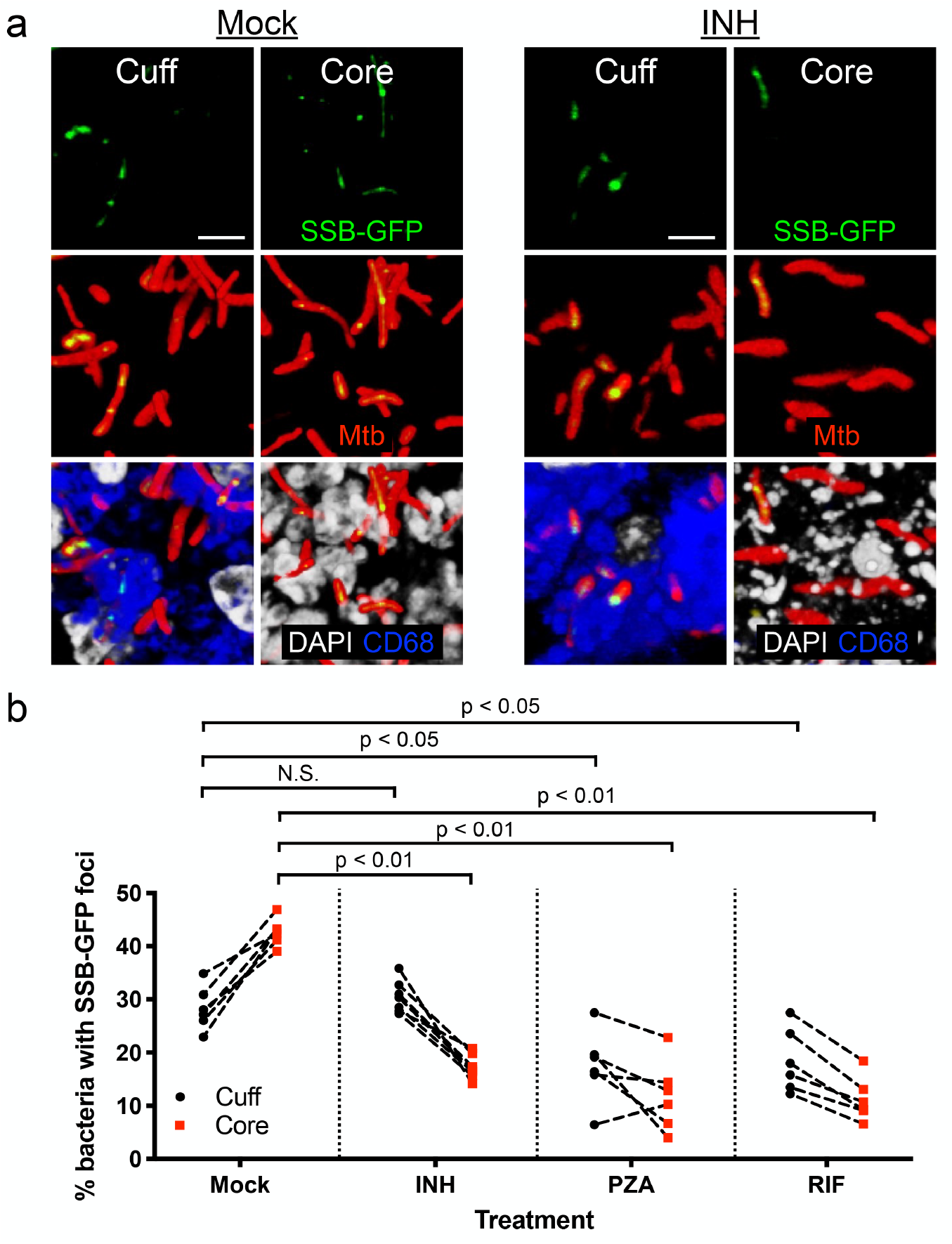
Impact of bacterial sublocation within caseous necrotic lesions on efficacy of first-line antitubercular drugs. (a) Representative 3D confocal images from the lesion cuff and core from a 6 week infection of C3HeB/FeJ mice with Erdman (SSB-GFP, *smyc’*::mCherry), followed by 2 weeks of mock or 10 mg/kg isoniazid (INH) treatment. All bacteria are marked in red (*smyc’*::mCherry), reporter signal is shown in green (SSB-GFP), nuclei are shown in grayscale (DAPI), and macrophages are shown in blue (CD68). For clarity of foci visualization, SSB-GFP signal is shown in extended focus, overlaid on the 3D images. Scale bar 5 μm. (b) shows the percentage of Mtb displaying SSB-GFP foci in each lesion sublocation for each quantified lesion, measured from multiple 3D confocal images for each set of 6 week infections followed by 2 weeks of mock, 10 mg/kg INH, 150 mg/kg pyrazinamide (PZA), or 10 mg/kg rifampicin (RIF) treatment. p-values were obtained with a Mann-Whitney statistical test. Sample details are as follows: mock treatment set – 6 different lesions from 5 mice; number of bacteria quantified in each lesion sublocation [cuff, core] was [641, 891], [481, 389], [408, 339], [318, 367], [447, 425], and [541, 453]. INH treatment set – 7 different lesions from 4 mice; number of bacteria quantified in each lesion sublocation [cuff, core] was [319, 371], [371, 467], [417, 385], [450, 403], [412, 419], [339, 311], and [368, 339]. PZA treatment set – 6 different lesions from 5 mice; number of bacteria quantified in each lesion sublocation [cuff, core] was [219, 210], [423, 626], [418, 543], [248, 419], [285, 366], and [341, 375]. RIF treatment set – 6 different lesions from 4 mice; number of bacteria quantified in each lesion sublocation [cuff, core] was [215, 297], [300, 151], [246, 275], [212, 122], [318, 378], and [367, 215].

In the case of PZA, treatment decreased the percentage of actively replicating Mtb in both lesion sublocations compared to the mock-treated control (Fig. 4b). While PZA has been shown to more efficacious at acidic versus neutral pH^14,15^, and the caseous necrotic core has a more neutral pH (Fig. 1)^8,9^, it is now also appreciated that PZA can have efficacy in conditions that do not involve non-acidic pH values^38,39^. Our observations here support the concept that PZA has efficacy even within the caseous necrotic core, suggesting that the *in vivo* effect of PZA extends beyond a dependency on local acidic pH conditions. Finally, while RIF treatment showed efficacy against the Mtb population present in both lesion sublocations, it decreased the percentage of Mtb with SSB-GFP foci in the caseous necrotic core to a greater extent (Fig. 4b). RIF acts to inhibit Mtb transcription^40–42^, and our findings here are thus in accord with its mode of action.

Despite the consensus on the importance of heterogeneity on Mtb infection progression and treatment outcome, population-level readouts such as bacterial load and host cytokine levels continue to be the primary means by which these outcomes are measured. Our establishment here of an integrated imaging approach for *in situ* tissue analysis presents a method to overcome the significant hurdle of the single bacterium resolution required for analysis of Mtb *in vivo* infection heterogeneity. We delineate pH and Cl^−^as two facets of the microenvironment that exhibit intra-lesion heterogeneity during Mtb infection, findings that expand on previous studies reporting on the neutral pH of liquefying caseum^8,9^, by providing single bacterium resolution analysis in both the lesion cuff and core sublocations, and revealing that heterogeneity in pH and [Cl^−^] is additionally present even within a lesion sublocation. Pimonidazole-based histological labeling has indicated the presence of hypoxia in the cuff of caseous necrotic lesions^22,43^, and we anticipate that other vital environmental signals such as nitric oxide are likely to also exhibit intra-lesion heterogeneity. Critically, our data directly demonstrate non-uniformity in bacterial replication status and transcriptional/translational activity within a single lesion, and reveal how these heterogeneous aspects are specifically linked to intra-lesion location. These differences in local environment and Mtb replication and activity further strikingly correlate to spatial differences in drug efficacy that exists even within a single caseous necrotic lesion. This work sets a foundational experimental method and framework for interrogation of (i) the heterogeneity in diverse aspects of Mtb infection biology within and between lesions in a single host, via the use of various reporter Mtb strains^33,44–47^, (ii) how drug treatment efficacy may differ in different sublocations within the lung/lesions and/or affect the local environment experienced by the bacterium, and (iii) how targeting of Mtb response to the local environment may change the extent of heterogeneity observed and thereby alter treatment success. We propose that future such single bacterium level studies in the context of intact tissue architecture, perturbing either regulators of bacterial environmental sensing and response or testing the effect of various therapeutic combinations, will build on the groundwork laid here and provide critical mechanistic insight into what drives infection heterogeneity, and how such non-uniformity impacts our ability to successfully treat Mtb infection.

## METHODS

### Mtb strains and culture

Reporter Mtb strains (*smyc’*::mCherry; *rv2390c’*::GFP, *smyc’*::mCherry; SSB-GFP, *smyc’*::mCherry) used for measuring differences in local microenvironment and replication status were in the Erdman background and have been previously described^12,23^. The P_606_’::mKO, *smyc’*::mCherry was as previously described^30^, except that the construct was placed in the background of the pDE43-MCK backbone, which is an integrating vector^48^. Bacteria were cultured in standing T25 flasks with filter caps, in 7H9 Middlebrook medium supplemented with OADC, 0.05% Tween 80, and 50 μg/ml hygromycin B or 25 μg/ml kanamycin as needed, buffered at pH 7.0 with 100 mM MOPS. Preparation of Mtb stocks for mice infection were as previously described^23^.

### Ethics statement

All animal procedures followed standards set by the National Institutes of Health “Guide for the Care and Use of Laboratory Animals”. Animal protocols were reviewed and approved by the Institutional Animal Care and Use Committee at Tufts University (#B2019-10), in accordance with the Association for Assessment and Accreditation of Laboratory Animal Care, the US Department of Agriculture, and the US Public Health Service guidelines. Light anesthesia during infection and oral gavage administration of drugs was via exposure to 2% isoflurane delivered by a vaporizer system. Euthanasia utilized carbon dioxide gas with regulated flow, consistent with American Veterinary Medical Association guidelines.

### Mouse Mtb infections

6-8 week old female C3HeB/FeJ wild type mice (Jackson Laboratory, Bar Harbor, ME) were intranasally infected with 10^3^ colony forming units of appropriate reporter Mtb strain in 35 μl of phosphate-buffered saline (PBS) containing 0.05% Tween 80, under light anesthesia with 2% isoflurane. Mice were sacrificed at 2, 6, 7, or 8 weeks post-infection, and the lungs fixed overnight in 4% paraformaldehyde (PFA) in PBS, before transfer and storage in PBS prior to analysis. For the P_606_’::mKO, *smyc’*::mCherry reporter strain infection, infections were allowed to establish for one or six weeks, before provision of the mice with water containing 5% sucrose ± 1 mg/ml doxycycline, with one additional water change during the one week treatment period. For the drug treatment infections, C3HeB/FeJ wild type mice were infected with the SSB-GFP, *smyc’*::mCherry reporter and infection allowed to establish for six weeks, before commencement of treatment with 10 mg/kg isoniazid (INH), 150 mg/kg pyrazinamide (PZA), or 10 mg/kg rifampicin (RIF) via oral gavage five times a week for 2 weeks (in a 200 μl volume). Control infected mice were mock-treated with sterile water. All drugs were prepared weekly, and all oral gavage treatments were carried out under light anesthesia with 2% isoflurane. INH and PZA working solutions were prepared directly in sterile water. RIF working solution was prepared by diluting 50 mg/ml RIF stock in DMSO to 10 mg/kg RIF + 5% DMSO in sterile water.

### Confocal immunofluorescence microscopy

Fixed lung lobes were embedded in 4% agarose in PBS and 250 μm sections obtained with a Leica VT1000S vibratome^49^. Staining of tissue was essentially as previously described^12,23,49^ – lung sections were blocked and permeabilized in PBS + 3% BSA + 0.1% Triton X-100 (“blocking buffer”) for 1 hour at room temperature, before incubation with primary antibodies overnight at room temperature (all steps on a nutator). The next morning, samples were washed 3 × 5 minutes with blocking buffer, then incubated with secondary antibodies at room temperature for 2 hours (all steps on a nutator). Samples were washed 3 × 5 minutes with blocking buffer again, and mounted with Vectashield mounting medium (Vector labs, Burlingame, CA). Rabbit anti-collagen I (Novus Biologicals, Centennial, CO, catalog #NB600-408) was used at 1:250, and Alexa Fluor 514 goat anti-rabbit (Invitrogen, Carlsbad, CA, catalog #A31558) used at 1:200 for secondary detection. Rat anti-CD68 (Bio-Rad, Hercules, CA, catalog #MCA1957) and rat anti-Ly6G (BD Biosciences, San Jose, CA, catalog #551459) were each used at 1:100, and Alexa Fluor 647 goat anti-rat (Invitrogen, catalog #A21247) used at 1:100 for secondary detection. Nuclei were visualized with DAPI (1:500; Invitrogen, catalog #D3571). Samples were imaged on a Leica SP8 spectral confocal microscope.

For broad xy-plane imaging, the Leica LAS X Navigator module was used to obtain multiple overlapping images that were then automatically merged together. High resolution images for reporter quantification were 10 μm in depth, reconstructed into 3D using Volocity software (Quorum Technologies, Ontario, Canada) from images taken at 0.5 μm z-steps. Quantification of reporter signal was carried out essentially as previously described using Volocity software^12,23,49^. In brief, for the *rv2390c’*::GFP reporter, the volume of each bacterium was measured via the mCherry channel, and the corresponding total GFP signal for that given object (bacterium) simultaneously measured. The settings for the GFP channel were maintained across samples to allow for comparison of values. Statistical analysis was performed using a Mann-Whitney test for comparison of reporter signal in bacteria present in the cuff versus core region in each lesion. A multiple t-test with a Holm-Sidak correction was used for statistical analysis of the binned data of reporter signal in the cuff versus core across all lesions. The P_606_’::mKO, *smyc’*::mCherry reporter strain was quantified in the same manner as for the *rv2390c’*::GFP reporter. For the SSB-GFP reporter, individual bacteria were identified via the mCherry channel and the number of bacteria with SSB-GFP puncta determined. Numbers of bacteria quantified in each case are indicated in the figure legends. Statistical analysis was performed using a Wilcoxon matched-pairs signed rank test for comparison of lesion cuff versus core values, and with a Mann-Whitney statistical test for comparing drug to mock treatment in a lesion sublocation.

## DATA AVAILABILITY

All relevant data are within the paper and its supplementary files.

## ACKNOWLEDGEMENTS

We thank Yuzo Kevorkian and Alwyn Ecker for excellent technical assistance, members of the Tan lab for helpful discussion, and Joan Mecsas for critical reading of the manuscript. This work was supported by grants R21 AI137759 and R01 AI143768 to ST from the National Institutes of Health. RCL was supported in part by training grant T32 AI007422 to Ralph R. Isberg from the National Institutes of Health. Confocal imaging was carried out in the Imaging Core Facility of the Tufts University Center for Neuroscience Research, supported by grant P30 NS047243 to F. Rob Jackson from the National Institutes of Health. The funders had no role in study design, data collection and analysis, decision to publish, or preparation of the manuscript.

