## Supplemental figures for "Spatial relationships of intra-lesion heterogeneity in *Mycobacterium tuberculosis* microenvironment, replication status, and drug efficacy"

**Supplementary Figure 1**

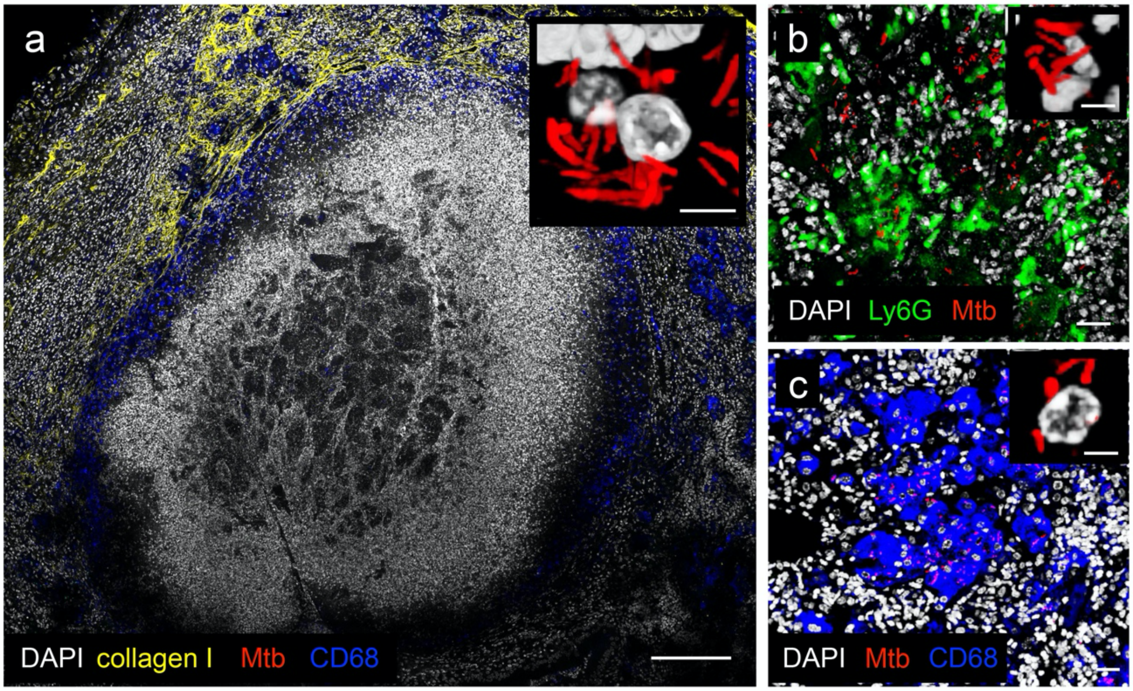


**Fig. S1. Visualizing Mtb infection at the single bacterium level within the context of intact tissue architecture.** Confocal image from 6 week infection of C3HeB/FeJ mice with Erdman (*smyc'*::mCherry) of (a) type I caseous necrotic lesion (~6 x 5 tiled image), (b) type II neutrophil-dominant lesion (~3 x 3 tiled image), and (c) type III macrophage-dominant lesion (~3 x 3 tiled image). All bacteria are marked in red (*smyc'*::mCherry) and nuclei shown in grayscale (DAPI) for all panels. Collagen I is shown in yellow in (a), and macrophages are shown in blue (CD68) in both (a) and (c). Neutrophils are shown in green (Ly6G) in (b). Insets show 3D confocal images of a targeted region from each respective panel, demonstrating single cell-resolution visualization of Mtb. Scale bar 200 μm in (a), 20 μm in (b) and (c), and 5 μm in insets.

**Supplementary Figure 2**

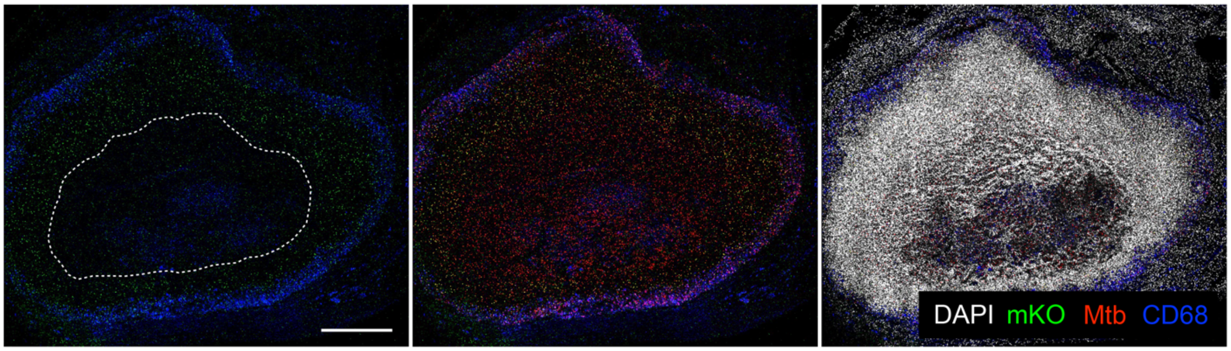


**Fig. S2. Inhibition of doxycycline penetration into the very central region of caseous necrotic lesions.** Overview confocal image (~11 x 9 tiled image) from a 6 week infection of C3HeB/FeJ mice with Erdman ($P_{606}$'::mKO-tetON, *smyc'*::mCherry), followed by 1 week of exposure to drinking water + 1 mg/ml doxycycline. All bacteria are marked in red (*smyc'*::mCherry), reporter signal is shown in green ($P_{606}$'::mKO-tetON), nuclei are shown in grayscale (DAPI), and macrophages are shown in blue (CD68). Scale bar 500 µm.
